# Spatial and Temporal Analysis of the Stomach and Small Intestinal Microbiota in Fasted Healthy Humans

**DOI:** 10.1101/450221

**Authors:** Anna M. Seekatz, Matthew K. Schnizlein, Mark J. Koenigsknecht, Jason R. Baker, William L. Hasler, Barry E. Bleske, Vincent B. Young, Duxin Sun

## Abstract

Although the microbiota in the proximal gastrointestinal (GI) tract has been implicated in health and disease, much of these microbes remains understudied compared to the distal GI tract. This study characterized the microbiota across multiple proximal GI sites over time in healthy individuals.

As part of a study of the pharmacokinetics of oral mesalamine administration, healthy, fasted volunteers (N=8; 10 observation periods total) were orally intubated with a four-lumen catheter with multiple aspiration ports. Samples were taken from stomach, duodenal, and multiple jejunal sites, sampling hourly (≤7 hours) to measure mesalamine (administered at t=0), pH, and 16S rRNA gene-based composition.

We observed a predominance of Firmicutes across proximal GI sites, with significant variation compared to stool. The microbiota was more similar within individuals over time than between subjects, with the fecal microbiota being unique from that of the small intestine. The stomach and duodenal microbiota displayed highest intra-individual variability compared to jejunal sites, which were more stable across time. We observed significant correlations in the duodenal microbial composition with changes in pH; linear mixed models identified positive correlations with multiple *Streptococcus* operational taxonomic units (OTU) and negative correlations with multiple *Prevotella and Pasteurellaceae* OTUs. Few OTUs correlated with mesalamine concentration.

The stomach and duodenal microbiota exhibited greater compositional dynamics compared to the jejunum. Short-term fluctuations in the duodenal microbiota was correlated with pH. Given the unique characteristics and dynamics of the proximal GI tract microbiota, it is important to consider these local environments in health and disease states.

## INTRODUCTION

The microbiota of the proximal gastrointestinal tract in humans represent an understudied yet highly relevant microbial community.^1^ Physiological processes such as gastric emptying, bile acid secretion, and the transit of food can influence the proximal gastrointestinal (GI) tract and disease development.^2-5^ However, our current understanding of how the processes and microbiota in different regions of the proximal GI tract relate to health and disease remains limited compared to other areas of the GI tract.

Much of our knowledge about the involvement of the human GI microbiota in health and disease has relied on fecal sampling, a non-invasive sampling method that is largely representative of the large intestine.^6,7^ Although it is known that the microbiota across the GI tract varies in composition and density,^8-10^ studying the microbiota at these sites is difficult, limiting our knowledge to invasive procedures, specific patient populations, or single time points.^1^ Analyses of mucosal samples from autopsies, endoscopies, and colonoscopies have revealed that Streptococci and Lactobacilli, both members of the oral and esophageal microbiota, are abundant members of the jejunal and ileal microbiota.^11-17^ Studies using naso-ileal catheters and ileostoma effluent, which allow collection over time, have supported these conclusions and revealed that the small intestinal microbiota is highly dynamic over short time courses, likely reflective of physiological processes at the stomach-small intestine interface.^18-21^

Understanding how these microbiota along the GI tract are related is of physiological relevance, particularly in relation to intestinal homeostasis and disease. Recent evidence suggests that the drug mesalamine, designed to reach high concentrations in the GI tract as treatment for irritable bowel disease (IBD), may directly target the microbiota in addition to host effectors.^22,23^ It is possible that some of the effectiveness of mesalamine treatment for IBD is mediated by the microbiota, potentiating the need to characterize these microbial communities to a fuller extent in the context of mesalamine administration.

This study investigated the bacterial composition across the intact upper GI tract in the same healthy, fasted adults over time. We used a multi-lumen tube designed to sample multiple sites along the upper GI tract. As part of a previously published study aimed at measuring mesalamine dissolution, subjects were given a dose of mesalamine and the proximal GI tract lumen was sampled over time.^24^ We used these samples to 1) characterize and compare microbial community dynamics over time at multiple upper GI sites within an individual and 2) identify how environmental factors, such as pH and the acute effect of mesalamine, shaped the microbiota. To the best of our knowledge, this is the first study to characterize the luminal microbiota across multiple upper GI sites over time within the same individual.

## METHODS

### Study recruitment

Healthy individuals (age 18-55) were included who were free of medications for the past two weeks, passed routine health screening, had a BMI 18.5-35, and had no significant clinical illness within three weeks. Health screening included a review of medical history and a physical examination (checking vital signs, electrocardiography, and clinical laboratory tests) described in Yu et al.^24^

### Catheter design and sterilization

A customized multi-channel catheter was constructed by Arndorfer Inc. (Greendale, WI), consisting of independent aspiration ports located 50 cm apart. The catheter had a channel to fit a (0.035 in x 450 cm) guidewire (Boston Scientific, Marlborough, MA), a channel connected to a balloon that could be filled with 7 ml of water to assist tube placement, and an end that was weighted with 7.75 grams of tungsten. Each single-use catheter was sterilized according to guidelines set by the American Society for Gastrointestinal Endoscopy at the University of Michigan prior to insertion (Supplemental Methods).^25^

### Collection of GI fluid samples

The full details of catheter placement have been previously described.^24^ Briefly, catheter placement occurred approximately 12 hours before sample collection. The catheter was orally inserted into the GI tract with aspiration ports located in the stomach, duodenum, and the proximal, mid and distal jejunum, confirmed by fluoroscopy. Subjects were given a light liquid snack approximately 11 hours before sample collection and fasted overnight for 10 hours prior to sample collection. At 0 hours, a mesalamine formulation was administered to each subject (Table 1). Luminal GI fluid samples (approximately 1.0 ml) were collected from up to four sites of the upper GI tract hourly up to 7 hours. Samples were collected by syringe, transferred to sterile tubes, and placed at −80°C until sample processing. A paired sample was collected to detect pH using a calibrated micro pH electrode (Thermo Scientific (Waltham, MA) Orion pH probe catalog no. 9810BN).

**Table 1:** Subject Recruitment. Selected metadata and sample collections for 10 admissions (subject M046 was admitted for three visits).

| Subject ID* | Mesalamine Formulation† | Age | BMI | Sex | Stomach | Duodenum | Jejunum |  |  | Stool | Total |
| --- | --- | --- | --- | --- | --- | --- | --- | --- | --- | --- | --- |
|  |  |  |  |  |  |  | Proximal | Mid | Distal |  |  |
| M046-A | Pentasa | 38 | 21.2 | M | 1 | 8 | - | 7 | - | 1 | 17 |
| M046-B | Apriso | 38 | 21.3 | M | - | 8 | - | 5 | 6 | - | 19 |
| M046-C | Lialda | 38 | 21.7 | M | 8 | 6 | - | 7 | - | 1 | 22 |
| M047 | Pentasa | 36 | 21.1 | M | - | 8 | 6 | - | - | - | 14 |
| M048 | Apriso | 51 | 34.3 | F | 5 | 7 | - | - | - | - | 12 |
| M053 | Apriso | 34 | 25.2 | F | 1 | - | 7 | 3 | - | - | 11 |
| M061 | Pentasa | 51 | 21.6 | M | 7 | 8 | - | - | - | - | 15 |
| M062 | Pentasa | 37 | 27.3 | M | 7 | 7 | - | - | - | 1 | 15 |
| M063 | Lialda | 26 | 28.6 | M | 7 | 5 | - | 5 | - | - | 17 |
| M064 | Lialda | 25 | 27.5 | F | 8 | 7 | - | - | - | - | 15 |
| **Summary** | **40% P, 30% A, 30% L** | **37 ±8.6** | **25 ±4.4** | **70% M** | **44** | **64** | **13** | **27** | **6** | **3** | **157** |

\*All subjects were caucasian and none identified as hispanic/latinx.
†Pentasa = Immediate release in stomach acid; Apriso = Extended release at pH > 6; Lialda = Extended release at pH > 7

### DNA extraction and Illumina MiSeq sequencing

The detailed protocol for DNA extraction and Illumina MiSeq sequencing was followed as previously described with modifications (Supplemental Methods).^26^ Briefly, 0.2 ml of GI fluid or 20 mg of stool was used for DNA isolation using a Qiagen (Germantown, MD) MagAttract Powermag microbiome DNA isolation kit (catalog no. 27500-4-EP). Barcoded dual-index primers specific to the V4 region of the 16S rRNA gene were used to amplify the DNA,^27^ using a “touchdown PCR” protocol (Supplemental Methods). Multiple negative controls were run parallel to each PCR reaction. PCR reactions were normalized, pooled and quantified.^28^ Libraries were prepared and sequenced using the 500 cycle MiSeq V2 Reagent kit (Illumina, San Diego, CA, catalog no. MS-102-2003). Raw FASTQ files, including those for negative controls, were deposited in the Sequence Read Archive database (BioProjectID: PRJNA495320; BioSampleIDs: SAMN10224451-SAMN10224634).

### Data processing and microbiota analysis

Analysis of the V4 region of the 16S rRNA gene was done using mothur (v1.39.3).^27,29^ Full methods, including detailed processing steps, raw processed data, and code for each analysis, are described in: https://github.com/aseekatz/SI_mesalamine. Briefly, following assembly, quality filtering, and trimming, reads were aligned to the SILVA 16S rRNA sequence database (v128).^30^ Chimeric sequences were removed using UCHIME.^31^ Prior to analysis, both mock and negative control samples (water) were assessed for potential contamination; samples with < 2500 sequences were excluded (Table S1). Sequences were binned into operational taxonomic units (OTUs), 97% similarity, using the opticlust algorithm.^32^ The Ribosomal Database Project (v16) was used to classify OTUs or sequences directly for compositional analyses (> 80% confidence score).^33^ Alpha and beta diversity measures (inverse Simpson index; the Yue&Clayton dissimilarity index, *θ*_*YC*_)^34^ were calculated from unfiltered OTU data. Basic R commands were used to visualize results, calculate % OTUs shared between samples, and conduct statistics, using packages plyr, dplyr, gplots, tidyr, and tidyverse. The nonparametric Kruskal-Wallis test, using Dunn’s test for multiple comparisons and adjusting *p*-values with the Benjamini-Hochberg method when indicated, was used for multi-group comparisons. The R packages lme4^35^ and lmerTest^36^ were used for mixed linear models between OTU relative abundance (filtered to include OTUs present in at least half of samples collected from a subject, per site) and pH or mesalamine.

## RESULTS

### Study population

Using a multi-channel catheter with multiple aspiration points,^24^ samples collected from the upper GI tract of 8 healthy subjects during 10 different study visits were processed for 16S microbial community analysis (Supplemental Methods, Table 1, Table S1). Samples were collected hourly over the course of 7 hours primarily from the proximal GI tract in the following possible locations: the stomach (n=44), duodenum (n=64), proximal/mid/distal jejunum (n=46), and stool (n=3). At the beginning of the study, subjects were given one form of mesalamine (Table 1). One of the seven subjects was studied three times over the course of 10 months; for most analyses, each study visit from this subject was considered independently.

### The proximal GI microbiota is dominated by Firmicutes and is distinct from the fecal microbiota

Analysis of the relative abundances of 16S rRNA-encoding genes from the GI tract across all timepoints and individuals demonstrated that the small intestinal microbiota was compositionally unique compared to stool (Fig. 1A). At all four sites in the proximal GI tract, Firmicutes composed the most abundant phyla (i.e. *Streptococcus, Veillonella*, and *Gemella* sp.). Higher levels of Bacteroidetes species (*Prevotella*) were detected in the stomach and duodenum. Proteobacteria and Actinobacteria predominated the remainder of the community at all sites. Diversity of the microbiota (inverse Simpson index) was decreased in sites of the upper GI tract compared to stool, which were enriched in Firmicutes (*Blautia*, Ruminococcaceae sp., and *Faecalibacterium*) and depleted in Bacteroidetes in these individuals (n=3) (Fig. 1B).

**Figure 1:**
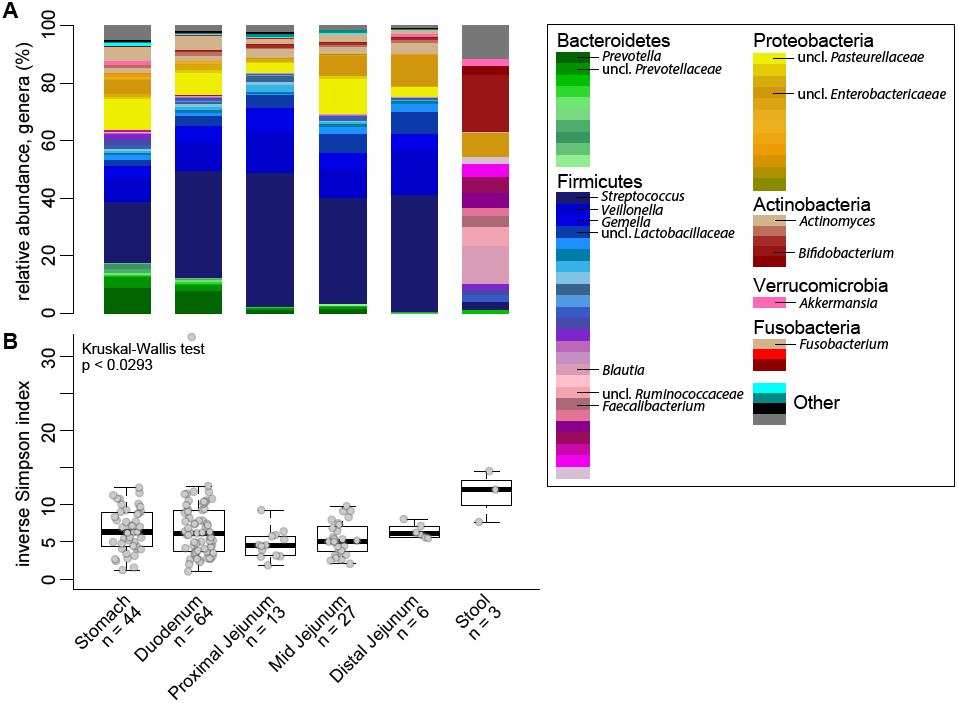
Bacterial community relative abundance and diversity in the upper GI tract. **A)** The mean relative abundance of genera at each GI site (sample *n* indicated). **B)** Boxplots of the inverse Simpson Index measuring community diversity across the GI tract (median, with first and third inter-quartile ranges). Statistical analysis: Kruskal-Wallis test (ns).

### The proximal GI microbiota is individualized and variable over time

To compare the microbiota across the proximal GI tract within and across individuals, we assessed pairwise community dissimilarity using the Yue&Clayton dissimilarity index, *θ*_*YC*_, which takes into account relative abundance of OTU compositional data. Both across (inter-individual) and within (intra-individual) subjects, stool was highly dissimilar to any proximal GI site (Figure 2A, 2B). Across proximal GI sites, subjects were more similar to their own samples than samples across other individuals (Figure 2A-D). The stomach microbiota was highly dissimilar across individuals compared to the duodenum or any part of the jejunum, which exhibited the least amount of dissimilarity (Figure 2C). A similar degree of dissimilarity was observed within an individual in the stomach, duodenum, and combined parts of the jejunum (Figure 2D).

**Figure 2:**
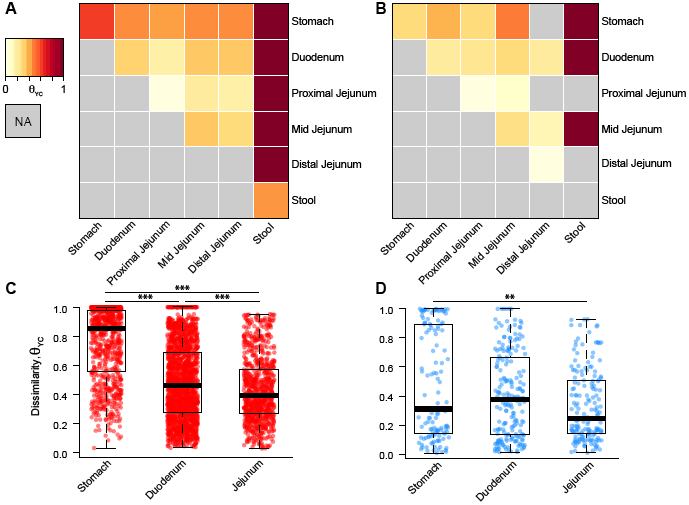
Dissimilarity of the proximal GI tract within and across individuals. Heatmap of the Yue&Clayton dissimilarity index, *θ*_*YC*_, comparing different proximal GI sites and stool A) across individuals (inter-individual pairwise comparisons) and C) within individuals (intra-individual pairwise comparisons). C) Inter-individual and D) intra-individual dissimilarity in the stomach, duodenum, and jejunum (sites combined). Statistical analysis: Kruskal-Wallis test (will add p values to graph). Statistical analysis: Dunn’s test for multiple comparisons with a Benjamini-Hochberg p-value adjustment (*p < 0.01; **p < 0.001; ***p < 0.0001).

Using a dissimilarity measure such as *θ*_*YC*_ allows us to assess stability based on changes in the relative abundance of OTUs. It is possible that certain GI sites fluctuate more in total OTUs. To measure whether any site had a higher rate of flux in their community, i.e. a higher rate of OTU turnover, we calculated the % OTUs detected at a given timepoint from the total number of OTUs detected within that individual at a given site. We observed that for each proximal GI site, a mean of 36.6% of the OTUs ever detected in that subject at a given site (mean number of total OTUs ever detected per subject per site = 135; range 78-212) were detectable at a given timepoint (Figure 3A). Similarly, we calculated the number of OTUs that were consistently present in all samples collected at that site within an individual (mean number of consistently detected OTUs per subject per site = 14.1; range 2-45). Overall, only 28.7% of the total OTUs ever detected at a given time point within an individual at a given site were represented by these consistently prevalent OTUs (Figure 3B). However, these prevalent OTUs explained an average of 72.0% of the relative abundance observed in the samples (Figure 3C). Of all sites, the relative abundance explained by the individual’s most prevalent OTUs in the stomach was lowest, followed by the duodenum, suggesting more variation at these sites compared to the jejunum (Kruskal-Wallis, p < 0.05).

**Figure 3:**
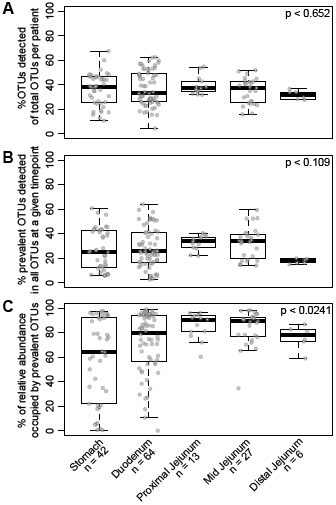
Fluctuations in prevalent OTUs observed within an individual across the proximal GI tract. **A)** Boxplots of the percentage of OTUs detected in a given sample out of all OTUs detected (all OTUs possible for that individual) at a subject-site. **B)** Boxplots of the percentage of OTUs that were consistently detected at a subject-site out of the total OTUs detected in a given sample. **C)** The percent of relative abundance explained by prevalent OTUs at a subject-site in a given sample. Statistical analysis: Kruskal-Wallis test.

One subject (M046) returned three times over the course of 10 months, allowing us to compare long-term changes. Across the sites that were sampled during multiple visits (the duodenum and mid-jejunum), prevalent OTUs were still detected during all three visits, explaining 74.4% and 66.1% OTUs in the duodenum and mid-jejunum, respectively (Fig. S1).

### Large fluctuations in the duodenal microbiota are associated with pH but not mesalamine

We next investigated how these compositional trends changed over time across the subjects. We focused on the duodenum and stomach since these sites were highly sampled across and within individuals and demonstrated variable pH. In the duodenum, we observed large fluctuations in genus-level composition across hourly timepoints within individuals (Figure 4, Figure S2, S3). Specifically, the relative abundance of *Streptococcus, Prevotella*, and an unclassified Pasteurellaceae species fluctuated in all individuals. We hypothesized that these fluctuations could be driven by mesalamine, administered in different forms to each subject at study onset. However, no visible pattern was observed with mesalamine levels. Interestingly, we observed that these compositional changes tracked with pH fluctuations (Figure 4). These patterns were less apparent in the stomach, where individuals displayed variable dynamics and highly individualized compositional patterns independent of mesalamine levels or pH, or in the jejunum of the subject with three different admissions, where pH fluctuated less (Figure S1, S2).

**Figure 4:**
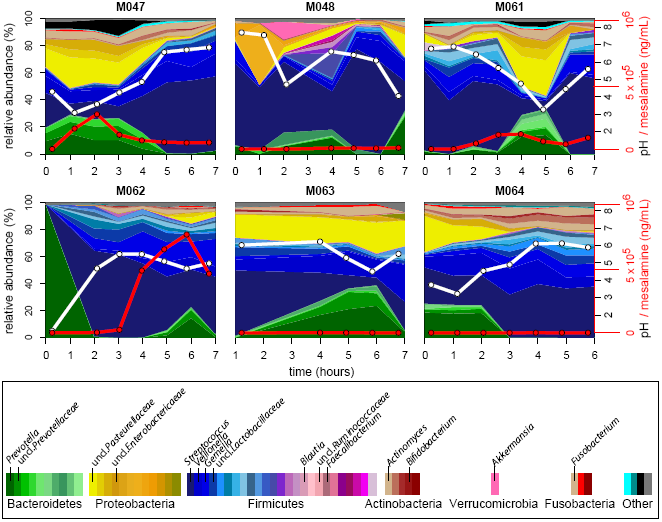
Longitudinal compositional dynamics, mesalamine levels, and pH in the duodenum. Streamplots of genus-level composition over time in the duodenum of six individuals (%, left y-axis; genera coded in legend). White lines indicate pH measurements (black y-axis labels on right) and red lines indicate mesalamine concentration (red *y-axis* labels on right).

To identify whether any singular OTUs correlated with changes in pH, we applied a generalized linear mixed model approach that takes into account subject-specificity.^37-39^ Within duodenal samples (n=56), we observed 15 OTUs that significantly correlated with pH changes. Linear regression of pH and relative abundance of these OTUs was significant across all samples (Figure 5; Table S2). Of the negatively correlated OTUs, six OTUs were classified as Bacteroidetes, mainly *Prevotella*, and two OTUs were classified as Pasteurellaceae sp. (Proteobacteria). The majority of the OTUs that were positively correlated with pH were Firmicutes, mainly *Streptococcus*, alongside an *Actinomyces* OTU (Actinobacteria). Only one OTU in the duodenum was significantly correlated to mesalamine (Table S2). We identified 17 OTUs that correlated with pH or mesalamine in the stomach; however, these were not representative at all sites (Table S2).

**Figure 5:**
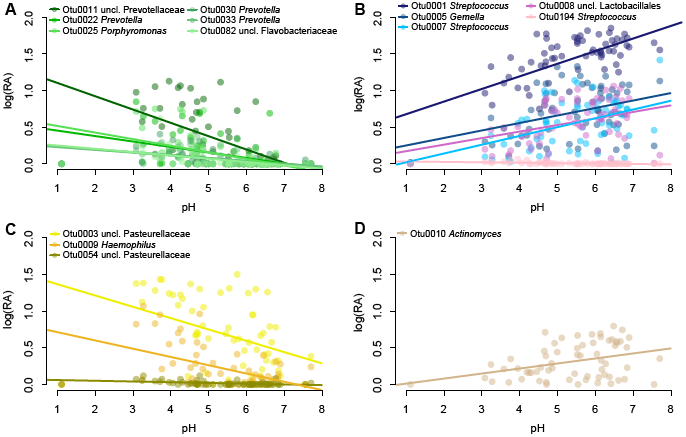
Relative abundance of significant OTUs vs. pH. Log relative abundance (log(RA)) as a function of pH of OTUs found to be significantly correlated with pH using linear mixed models (all samples with measurable pH). Lines represent linear fit per OTU. OTUs classified as **A)** Firmicutes, **B)** Bacteroidetes, **C)** Proteobacteria, and **D)** Actinobacteria are depicted (genus-level OTU classification coded by legends).

## DISCUSSION

Our results demonstrate that the microbial communities inhabiting the GI tract are distinct and dynamic across different sites within the proximal GI tract. Our sampling procedure provided us with an opportunity to longitudinally characterize such microbial populations in conjunction with the administration of a commonly used drug, mesalamine. We observed high stability of the microbiota in the jejunum compared to the stomach or duodenum, indicating that the indigenous microbiota residing in more proximal regions of the GI tract may experience greater changes. While we did not observe strong correlations between mesalamine concentration and particular microbiota members at any site, we did observe a strong correlation between the microbiota composition and pH, particularly in the duodenum.

In this report, we describe the use of a multi-lumen catheter design with unique aspiration ports that enabled sampling of small intestinal content over the course of seven hours.^24^ Many studies aimed at investigating the microbiota of the proximal GI have overcome sampling difficulty in this region by using ileostoma effluent, samples from newly deceased individuals, or naso-ileal tubes. Although easy to access, ileostoma effluent does not fully recapitulate the distal small intestine, as it more closely resembles the colon than the small intestine due to increased oxygen concentrations near the stoma.^40-43^ Single lumen naso-ileal tubes are unable to sample multiple sites simultaneously.^18,20,21,44^ GI fluid collected with our methodology was sufficient for determining mesalamine concentration, assaying fluid pH, and isolating microbial DNA across time and GI sites, which has not been previously described.^24^

Our results support previous observations that the small intestine is dynamic with higher inter-individual than intra-individual variability.^18,21,45^ However, the mid-distal small intestine also contains a resilient microbial community composed of several highly abundant OTUs. This resilience is demonstrated by the shift from an altered to a normal ileal microbiota following the resolution of an ileostoma.^46^ This mirrors the colonic microbiota, which also has a small community which is stable over long periods of time.^42,47,48^

This and other studies have shown that the jejunal and proximal ileal microbiota are distinct from the colonic microbiota.^10,49^ Despite changes in overall community structure and an overall decrease in microbial diversity across the stomach and small intestine compared stool, many of the same organisms commonly observed in stool were also present in the upper GI tract, albeit at very different abundances.^10^ Interestingly, colonic resection and ileal pouch-anal anastomosis has been shown to shift the terminal ileum microbiota to a state similar to the colon, suggesting that a colonic community structure can develop at these sites given the right conditions.^21,43,49-51^

Many of the abundant microbes observed in our study, *Streptococcus, Veillonella, Gemella*, and Pasteurellaceae species, are also common residents of the oral cavity, which reflects the proximity of these locations in the GI tract. Populations of Proteobacteria, such as Pasteurellaceae, have also been observed consistently in the small intestinal microbiota in other studies, particularly in patients with IBD.^14,52-54^ In our study, *Streptococcus* and *Veillonella* were correlated with pH in duodenal samples. It is possible that growth of these organisms drives a decrease in pH via metabolism of short-chain fatty acids, an observed functional capacity of these genera.^21,55^ Conversely, large fluctuations in environmental pH may select for genera like *Streptococcus*, which have evolved a variety of mechanisms to control pH intracellularly.^56-59^ In any case, our data suggests a relationship between microbial dynamics and environmental physiology of the duodenum, which is an important observation to consider when comparing this site across individuals.

We observed little association between mesalamine concentration and changes in microbial relative abundance in our cohort. Several studies have reported differences in the fecal microbiota of patients with or without IBD, in particular Crohn’s disease, which can affect the small intestine.^52^ Compositional shifts in the small intestine have been reported during IBD, specifically increased levels of Enterobacteriaceae species, such as *Enterococcus, Fusobacterium*, or *Haemophilus*.^14,53,54^ It has been hypothesized that mesalamine’s ability to reduce inflammation in patients with ulcerative colitis could be by altering the microbiota.^22,23^ While acute effects of mesalamine on the microbiota have not previously been reported, earlier work has demonstrated that mesalamine decreases bacterial polyphosphate accumulation and pathogen fitness, suggesting an influence on the microbiota.^23^ We did not observe strong correlations between mesalamine concentration and the microbiota here. However, our study was small, used different doses of mesalamine that may be metabolized differently across GI sites, and was conducted in healthy individuals.^24^ It is possible that mesalamine is less likely to impact the small intestinal microbiota, which historically has lower efficacy in treating active Crohn’s Disease, which manifests in the small intestine, compared to ulcerative colitis, which manifests in the large intestine.^22,60,61^ As indicated by the variability of mesalamine in the subjects in this study, the effects of mesalamine on the small intestinal microbiota may be highly individualized.^24,62-64^ Furthermore, individuals with disease may harbor a distinct microbiota that responds to mesalamine differently.

Despite the opportunity provided by our method to describe the microbiota across the GI tract, our study has some lingering questions. Movement by the subject during the study can result in movement of each sampling port, particularly between the distal stomach and antrum. This may explain the inconsistent pH values and severe fluctuations of the microbiota observed in the stomach. Similarly, the shorter length of the sampling device, as compared to a naso-ileal catheter, prevented reliable collection of fluid from the distal small intestine, limiting our sampling to the proximal region. We also were limited to three concurrent fecal samples, each of which was low in Bacteroidetes, a profile generally observed in individuals on low fat-high fiber, non-Western diets.^65^

The use of a novel catheter allowed us to assess the microbiota across several proximal GI sites overtime, representing a powerful clinical and/or investigative tool for studying the small intestinal microbiota. Future studies on the upper GI microbiota should collect concurrent oral swab/sputum and fecal samples to strengthen the ability to “track” microbial populations across the GI tract, potentiating our ability to correlate the microbiota from fecal sampling, a more convenient method to study the microbiota, to other sites of the GI tract.

## Declarations

## Acknowledgements

This research was funded by FDA grant HHSF223201000082C. Clinical samples collected with help from Michigan Institute for Clinical&Health Research (MICHR) NIH grant UL1TR000433. The authors would also like to thank the Host Microbiome Initiative and the Microbial Systems Molecular Biology Laboratory at the University of Michigan for their support with the 16s rRNA sequencing, and Krishna Rao and Rose Putler for their assistance with the modelling and statistical analyses. This research was funded by the FDA(HHSF223201000082C) and the NIH (5U01AI124255-03).

## Availability of data and material

SRA: BioProjectID: PRJNA495320; BioSampleIDs: SAMN10224451-SAMN10224634 GitHub: https://github.com/aseekatz/SI_mesalamine

## Authors’ contributions

MJK, DS, BEB, AMS, VBY, MKS - Conception or design of the work

MJK, BEB, WLH, JRB - Data collection

AMS, MJK, MKS - Data analysis and interpretation

MKS, AMS - Drafting the article

MKS, AMS, DS, VBY - Critical revision of the article

AMS, MKS, MJK, JRB, WLH, BEB, VBY, DS - Final approval of the version to be published

## Ethics approval and consent to participate

Samples collected in this study were part of clinical trial NCT01999400. The institutional review boards at the University of Michigan (IRBMED) and the Department of Health and Human Services, Food and Drug Administration (Research Involving Human Subjects Committee/RIHSC) both approved the study protocol. All subjects provided written informed consent in order to participate.

## Consent for publication

Not applicable

